# Selective Elimination of Osteosarcoma Cell Lines with Short Telomeres by ATR Inhibitors

**DOI:** 10.1101/2020.08.18.254664

**Authors:** Tomas Goncalves, Georgia Zoumpoulidou, Carlos Alvarez-Mendoza, Caterina Mancusi, Laura C. Collopy, Sandra J. Strauss, Sibylle Mittnacht, Kazunori Tomita

## Abstract

To avoid replicative senescence or telomere-induced apoptosis, cancers employ telomere maintenance mechanisms (TMMs) involving either the upregulation of telomerase or the acquisition of recombination-based alternative telomere lengthening (ALT). The choice of TMM may differentially influence cancer evolution and be exploitable in targeted therapies. Here, we examine TMMs in a panel of seventeen osteosarcoma-derived cell lines defining three separate groups according to TMM. Eight were ALT-positive, including the previously uncharacterised lines, KPD and LM7. ALT-negative cell lines were further classified into two groups according to their telomere length. HOS-MNNG, OHSN, SJSA-1, HAL, 143b and HOS displayed sub-normally short telomere length, while MG-63, MHM and HuO-3N1 displayed long telomeres. Importantly, sub-normally short telomeres were significantly associated with hypersensitivity to three different therapeutics targeting the ataxia telangiectasia and Rad3-related (ATR) kinase - AZD-6738/Ceralasertib, VE-822/Berzoserib and BAY-1895344 - compared to long telomeres, maintained via ALT or telomerase. Within 24 hours of ATR inhibition, cells with short but not long telomeres displayed chromosome bridges and underwent cell death, indicating a selective dependency on ATR for chromosome stability. Collectively, our work provides a resource to identify links between TMMs and drug sensitivity in osteosarcoma and indicates that telomere length predicts ATR-inhibitor sensitivity in cancer.

## Introduction

Telomeres are the protective ends of chromosomes, essential for all dividing cells [1]. While telomeres shorten with every chromosome replication cycle due to the end-replication problem, critically shortened telomeres elicit DNA damage checkpoint activation, leading to replicative senescence or apoptosis [2]. The vast majority of cancer cells maintain the ends of their chromosomes through a telomere maintenance mechanism (TMM) thereby avoiding senescence or death induced by critically shortened telomeres [3]. In about 90% of cancers, the reverse transcriptase enzyme telomerase, which replenishes telomeric DNA, is reactivated, permitting indefinite cell divisions [4, 5]. Only in exceptional circumstances, where cancer cells retain substantially long telomeres, TMMs may be bypassed [6–8].

Interestingly, the activity or expression levels of telomerase in cancers does not seem to correlate with the length of telomeres that are maintained [6, 9, 10]. A substantive fraction of telomerase-positive cancers display abnormally short telomeres compared to adjacent noncancerous tissue [6, 11–15]. Furthermore, shortness of telomeres in tumours is correlated with poor prognosis in many types of cancer [14–16]. Whereas the origin of short telomeres in tumours has not been clearly demonstrated, shortness or loss of telomeres is thought to originate from excessive divisions at the benign stages [1, 2, 17]. In telomerase-expressing normal cells, telomerase preferentially elongates the shortest telomeres to extend overall telomere length [18, 19]. However, such telomerase action appears to be compromised in cancer cells, resulting in retention of critically short telomeres and inefficient telomere lengthening [20–22]. The ability to tolerate and mechanisms to maintain these short telomeres by telomerase are thought to be unique to cancer [2, 21].

Telomerase activity has been targeted for many types of cancer. However, the cytotoxic effects of telomerase inhibition are not immediate as time is required to achieve telomere attrition and damage accumulation, with cancer cells able to propagate and evolve until their telomeres significantly shorten and become deprotected [7, 8, 23, 24].

In the absence of telomerase activity, some tumours gain the ability to undertake telomerase-independent, alternative maintenance of telomeres. Alternative lengthening of telomeres (ALT) is dependent on homologous recombination or homology-directed repair-based DNA replication [25, 26]. ALT-type TMM is often found in cancers of a mesenchymal origin, including central nervous system cancers, as well as in many sarcomas, particularly osteosarcoma, potentially explained by the fact that telomerase expression in cells of mesenchymal origin is more strictly regulated compared to cells of epithelial origin [5, 27–29]. To date, ALT activity has not been detected in normal human body cells, highlighting a unique cancer-selective therapeutic opportunity. However, despite this selectively cancer-associated feature, effective drugs that kill ALT-positive cancer cells are yet to be discovered.

Osteosarcoma is a rare cancer, commonly found in young adults. The overall five-year survival rate has not significantly improved over the past two decades and therapy-associated toxicities pose a significant problem in the treatment of this cancer [30, 31]. Therefore, new, less toxic strategies are needed. As children have significantly longer telomeres in their somatic cells [32], targeting the mechanisms of short telomere maintenance may pose unique opportunities for treatments that effectively eradicate cancer cells yet minimise normal tissue toxicity and side effects. To this end, we characterised TMM and telomere length within a panel of seventeen osteosarcoma cell lines [33] (Supplementary Table 1), sorting them into three separate categories based on mechanism of telomere lengthening and observed telomere length. We provide evidence of different telomere anatomies in telomerase-expressing ALT-negative osteosarcomas, evidenced by either overall long or short telomere length and report synthetic lethality of telomerase-expressing cells with short telomeres to clinical inhibitors of the Ataxia Telangiectasia and Rad3-Related (ATR) Ser/Thr kinase (ATRi). Our data highlight the use of telomere length as a biomarker to identify ATRi sensitivity in osteosarcoma and potentially other cancers.

## Results

### Categorisation of osteosarcoma cell lines by mode of telomere maintenance mechanism

Of the osteosarcoma cell lines in our panel, six (G292, HuO-9, CAL72, U2OS, NY and SAOS-2) have previously been characterised as ALT-positive [34, 35], while five (HOS-MNNG, SJSA-1, HOS, MG-63 and HuO-3N1) had previously been characterised as telomerase-positive and ALT-negative [35–39] (Table 1). However, TMM in a further six cell lines (OHSN, HAL, 143b, MHM, KPD and LM7) had not previously been characterised. Therefore, we first sought to assess the TMM used in these remaining lines alongside those previously assessed, to confirm the respective published results.

**Table 1.** Telomere status of the Osteosarcoma Cell Lines.

| Cell Line | TMM | qPCR T/S Ratio | Median Telomere Length (kb)* | CC Assay Level | TERC level** | TERT mRNA level** |
| --- | --- | --- | --- | --- | --- | --- |
| **HOS-MNNG** | ST | 0.813 | 3.0 | -0.006 | 19.80% | 56.54% |
| **OHSN** | ST | 0.837 | 3.2 | -0.011 | 23.17% | 107.37% |
| **SJSA-1** | ST | 0.840 | 3.2 | -0.002 | 6.36% | 695.03% |
| **HAL** | ST | 0.853 | 2.7 | 0.007 | 1.48% | 170.34% |
| **143b** | ST | 0.855 | 3.0 | 0.002 | 26.48% | 288.96% |
| **HOS** | ST | 0.902 | 3.6 | 0.001 | 21.03% | 323.13% |
| **MG-63** | LT | 0.966 | 5.8 | -0.003 | 11.91% | 58.47% |
| **MHM** | LT | 1.086 | 6.4 | 0.001 | 29.03% | 390.11% |
| **HuO-3N1** | LT | 1.129 | 6.7 | 0.029 | 35.41% | 7.69% |
| **G292** | ALT | 1.415 | 10.7 | 4.156 | 3.98% | 0.06% |
| **HuO-9** | ALT | 1.539 | 9.7 | 1.066 | 3.69% | 1.33% |
| **CAL72** | ALT | 2.853 | 10.2 | 5.422 | 5.30% | 2.22% |
| **U2OS** | ALT | 4.221 | 13.7 | 0.835 | 0.00% | 4.21% |
| **KPD** | ALT | 1.210 | 10.0 | 2.944 | 0.01% | 4.42% |
| **NY** | ALT | 1.552 | 9.9 | 0.651 | 0.77% | 14.54% |
| **SAOS-2** | ALT | 1.230 | 7.6 | 1.975 | 21.87% | 19.36% |
| **LM7** | ALT | 1.170 | 8.2 | 1.227 | 12.27% | 0.29% |

ST – ALT-negative, short telomere
LT – ALT-negative, long telomere
ALT – ALT-positive
\*calculated using Telometric software
\*\*relative to HEK293T expression, determined as expression fold change versus 7SK

**Table 2.** Area Under the Curve (AUC) Analysis of ATRi.

| **Cell Line** | **TMM** | **AZD-6738** | **VE-822** | **BAY-1895344** |
| --- | --- | --- | --- | --- |
| **HOS-MNNG** | ST | 181.8±1.5 | 140.5±2.2 | 61.2±0.8 |
| **OHSN** | ST | 213.7±9.3 | 169.7±12.0 | 47.5±1.1 |
| **SJSA-1** | ST | 363.0±1.3 | 256.4±1.8 | 205.6±8.2 |
| **HAL** | ST | 272.7±6.9 | 214.7±0 | 136.9±1.7 |
| **143b** | ST | 215.9±6.8 | 128.6 | 57.4 |
| **HOS** | ST | 220.4±32.4 | 176.8±9.1 | 50.4±15.8 |
| **MG-63** | LT | 294.2±1.2 | 241.8 | 125.4 |
| **MHM** | LT | 275.4±7.4 | 190.7±6.3 | 111.1±0.8 |
| **HuO-3N1** | LT | 334.0±18.7 | 270.4±30.3 | 185.4±26.8 |
| **G292** | ALT | 288.5±35.4 | 237.7±43.2 | 189.3±19.4 |
| **HuO-9** | ALT | 242.8±16.5 | 217.5 | 149.2 |
| **CAL72** | ALT | 226.6±0.7 | 146.6±0.4 | 77.9±4.4 |
| **U2OS** | ALT | 305.6±16.0 | 245.8±8.1 | 158.9±16.5 |
| **KPD** | ALT | 254.0±37.5 | 155.5 | 188.3 |
| **NY** | ALT | 278.0±30.0 | 245.2 | 201.5 |
| **SAOS-2** | ALT | 304.0±13.5 | 246.2 | 176.6 |
| **LM7** | ALT | 321.7±1.3 | 251.5±17.9 | 195.9±12.1 |

ST – ALT-negative, short telomere
LT – ALT-negative, long telomere
ALT – ALT-positive

ALT-positive cells are characterised by the presence of C-circles, a by-product of telomere recombination in ALT, consisting of partially double-stranded extra-chromosomal telomeric DNA circles. C-circles were detected through rolling circle amplification (RCA) followed by quantification of telomeric sequences using either dot blot hybridisation or monochrome-multiplex quantitative polymerase chain reaction (MM-qPCR) (Figure 1a and Supplementary Figure 1a). Another hallmark of ALT-based telomere maintenance is the co-localisation of telomere clusters with promyelocytic leukaemia (PML) bodies, forming structures termed ALT-associated PML bodies (APBs) [29]. The C-circle positive cell lines, along with three cell lines that were C-circle negative, were subjected to the APB assay (Figure 1b, c and Supplementary Figure 2). This combined analysis defined the previously uncharacterised lines KPD and LM7 as ALT-positive, along with previously reported ALT-positive lines G292, HuO-9, CAL72, U2OS, NY and SAOS-2. We also defined the previously uncharacterised lines OHSN, HAL, 143b and MHM as ALT-negative, along with the previously ALT-negative lines HOS-MNNG, SJSA-1, HOS, MG-63 and HuO-3N1 (Table 1).

**Figure 1.**
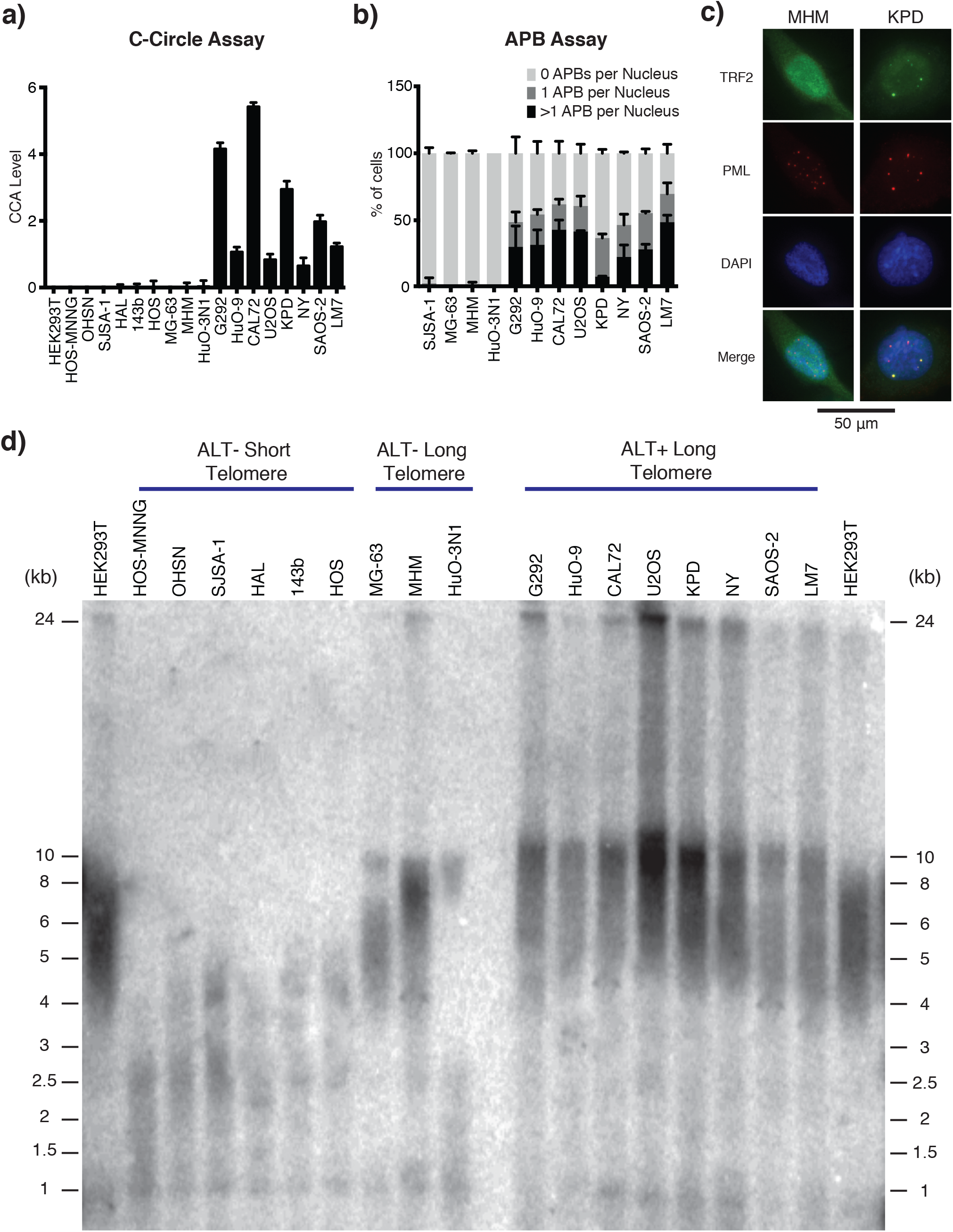
Characterisation of Telomere Status in Osteosarcoma Cell Lines. a) *C-Circle Assay (CCA) Level*. Graph shows the CCA level calculated by subtracting the phi-29 polymerase NTC by the phi-29 polymerase free control NTC (see methods). A sample is then considered ALT+ if its CCA level lower error bar is >0. Data is represented as a mean of three independent experiments. Error bars show standard deviation. Dot blot result of phi-29 polymerase is shown in Supplementary Figure 1a. b) *APB Assay for the osteosarcoma cell lines with long telomeres*. APBs were scored as a direct co-localisation between TRF2 and PML foci. The graph shows the mean percentage of cells containing 0, 1 or >1 APBs in their nuclei for three independently prepared samples of >50 cells. Error bars show standard deviation. c) *Representative immunofluorescence images showing the presence/absence of APBs in the KPD and MHM cell lines respectively.* TRF2 was stained green and PML bodies were stained red. Yellow foci (co-localisation of green and red foci) in the merge panel represent APBs. A 50-μm scale bar is shown. Representative images of all tested cell lines are shown in Supplementary Figure 2. d) *Telomere Southern blot of osteosarcoma cell lines*. Terminal restriction fragment length analysis of genomic DNA digested with *Hin*fI and *Rsa*I and hybridised with a telomeric TTAGGG probe. ALT-negative cell lines tend to have shorter telomeres than HEK293T, while ALT-positive cell lines tend to have longer telomeres than HEK293T, which are extremely heterogeneous in length. Digital analysis of telomere length is shown in Table 1.

The ALT pathway involves recombination-mediated robust amplification of telomeric DNA repeats through break-induced replication, generating heterogeneous telomeres that are significantly longer compared to those in telomerase-positive cancer cells. To determine the median length of telomeres in each of the various cell lines, we used telomere restriction fragment (TRF) assays where telomere length is determined by southern blotting (Figure 1d, Supplementary Figure 1b and Supplementary Table 3). Previous work proposed that ALT status may be predicted based on excessively long telomeres, with mean, variance and semi-interquartile range of telomere length distribution greater than 16 kb, 200 kb^2^ and 4 kb, respectively [29]. Consistent with this view we observed that all of the ALT-positive cell lines exhibited heterogenous telomere signals reaching over 24 kb in length. Three of the ALT-negative lines, MG-63, MHM and HuO-3N1, harboured telomere length comparable to the telomerase-positive noncancerous cell line HEK293T, serving as a control. Notably, the remaining six ALT-negative cell lines (HOS-MNNG, OHSN, SJSA, HAL, 143b and HOS) had significantly shorter telomeres and hence may represent situations of telomere maintenance present in cancers with sub-normally short telomeres.

To independently confirm the telomere length predictions based on TRF assessment, we used MM-qPCR, which determines telomeric repeat number relative to the repeat number of a reference gene [40]. In addition, MM-qPCR is suitable to determine telomere status in patient derived tumour samples [14, 41, 42]. To reduce errors caused by potential aneuploidy that may variably exist in the cancer lines, we used two different single copy number genes (SCGs) present on different chromosomes, *ALB* encoding albumin (4q13.3) and *HBB* encoding beta-globin (11q15.4). The ratio of telomere repeats to single-copy genes (T/S) were calculated (Supplementary Figure 1c). More uniform cycle threshold (Ct) values were obtained using *ALB*, suggesting a more reliable SCG for MM-qPCR analysis (Supplementary Figure 1d). Using the HEK293T cell line as a reference, the MM-qPCR based telomere length analysis correlates with the TRF-based analysis and clearly distinguished the cell lines into three categories (Supplementary Figure 1e,f).

Together, the above results put the cell lines into three clearly defined categories: 1) ALT-negative cell lines with short telomeres, 2) ALT-negative cell lines with long telomeres, and 3) ALT-positive cell lines with long telomeres (Table 1).

### Molecular profiling of osteosarcoma cell lines

To catalogue the molecular drivers involved in maintaining telomeres in the various osteosarcoma cell lines, we surveyed candidate molecular events known to be associated with different forms of telomere maintenance.

ALT is strongly associated with dysfunction of the alpha-thalassemia/mental retardation syndrome X-linked (ATRX) and the death domain-associated protein 6 (DAXX) chromatin remodelling complex [34]. In our panel, the ALT-positive cell lines, with the exception of G292 and KPD, lacked detectable ATRX expression (Figure 2a). Interestingly, the ALT-negative group of the osteosarcoma lines with short telomeres expressed remarkably high levels of ATRX while those with long telomeres had either low or undetectable level of ATRX. All cell lines expressed DAXX at the expected molecular weight, except G292, where DAXX is known to be fused to the kinesin family member KIFC3 [35]. The gene encoding SMARCAL1 (SWI/SNF-related Matrix-associated Actin-dependent Regulator of Chromatin subfamily A-Like protein 1) was recently found to be mutated in a number of ALT-positive glioblastomas [43]. In our osteosarcoma panel, only NY lacked SMARCAL1 expression, as was previously reported [35]. We did not find a loss of expression in the genes associated with ALT signature in the newly identified ALT-positive cell line KPD (Figure 2a). Further investigation will be required to elucidate the genetic basis leading to ALT activation in this cell line.

**Figure 2.**
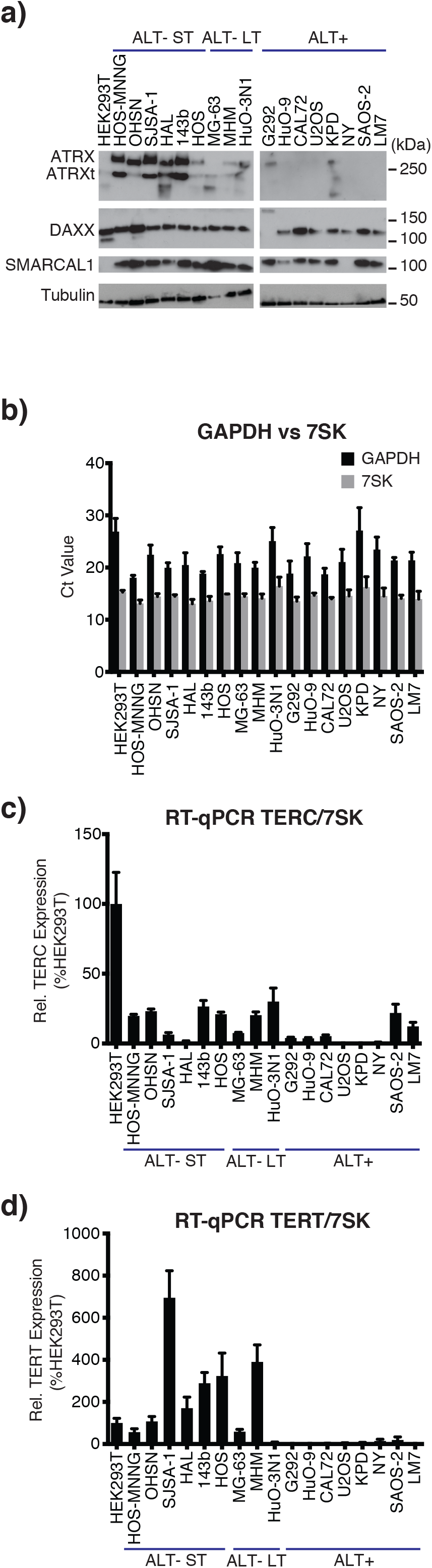
ALT-specific Mutations and Expression of Telomerase. a) *ATRX, DAXX and SMARCAL1 Western blot*. Western blot analysis of the osteosarcoma cell lines for ATRX, DAXX and SMARCAL1, three previously reported genes with mutations associated with ALT. G292 has a chimeric DAXX-KIFC3 fusion protein, as reported [35]. NY has no full length SMARCAL1 protein, as reported [35]. Alpha-tubulin was used as a loading control. ATRXt = truncated isoform of ATRX. b) *RT-qPCR: 7SK RNA is a more reliable reference than GAPDH mRNA*. Graph shows CT values of GAPDH (black) and 7SK (grey) from normalised mRNA in RT-qPCR. Data is represented as a mean of three independent experiments, each run in triplicate. Error bars show standard deviation. c) *RT-qPCR: TERC/7SK*. TERC was quantified using RT-qPCR and normalised first to 7SK and then to expression in the HEK293T cell line. There is no correlation between TERC expression and telomere length. Data represented as a mean of three independent experiments. Error bars show standard error of the mean. d) *RT-qPCR: TERT/7SK*. TERT mRNA was quantified using RT-qPCR and normalised first to 7SK and then to expression in the HEK293T cell line. There is no correlation between TERT expression and telomere length. Data represented as a mean of three independent experiments. Error bars show standard error of the mean.

Previous studies showed that the ALT-negative cell lines, HOS-MNNG, SJSA-1, 143b, HOS and MG-63 express telomerase [36–39, 44, 45]. In contrast, the ALT-positive cell lines, G292, HuO-9, Cal72, U2OS and NY, do not express telomerase while SAOS-2 expresses telomerase at low levels [37, 38, 45]. To confirm these published results and to determine telomerase status in the newly characterised osteosarcoma lines, we assessed the transcriptional levels of the telomerase subunits, telomerase reverse transcriptase (*TERT*) and its RNA component (*TERC*) using RT-qPCR.

Since cancer cells have variable transcription patterns even if of the same tissue of origin, we first investigated the expression level of two housekeeping genes, *GAPDH* and *7SK*. We found that *7SK* was a more reliable reference RNA as the standard deviation, standard error of the mean and coefficient of variation of the calculated Ct values across the cell lines were all smaller for *7SK* than for *GAPDH* (0.91 v 2.65, 0.21 v 0.63 and 6.28% v 12.27%, respectively), indicating that *7SK* has smaller relative variability (Figure 2b).

Using 7SK RNA as the reference, *TERC* and *TERT* mRNA expression were measured. In general ALT-negative cell lines expressed *TERC* at higher levels than the ALT-positive cell lines, with the exception of SAOS-2 and LM7 (Figure 2c). Most of the ALT-negative cell lines had *TERT* expression comparable to or greater than the HEK293T control cell line (Figure 2d). Exceptionally, *TERT* expression was notably low (7.7% of HEK293T) in the ALT-negative long telomere cell line HuO-3N1. In the ALT-positive cell lines, *TERT* expression level was consistently negligible/undetectable (Figure 2d). Overall, our analysis documents telomerase expression almost exclusively in ALT-negative osteosarcoma lines and confirms that there was no correlation between *TERC* or *TERT* expression and telomere length in these lines (Figure 2c, d).

### Short telomeres predict hypersensitivity to the ATR inhibitors in osteosarcoma

Our telomere length analysis categorised the ALT-negative osteosarcoma cell lines into two types: short and long telomere maintenance. We sought to identify therapeutic options that could synergise with short telomeres and hence may be beneficial in the treatment of osteosarcoma with this feature. Short telomeres are presumed to yield increased vulnerability to events that perturb telomere replication. The DNA damage/replication checkpoint kinase, ATR, is essential for facilitating replication fork progression [46], and counteracts telomere shortening in yeasts [47–49]. Importantly, ATR contributes to the maintenance of shortened telomeres in mice [19].

We therefore assessed the response of the various osteosarcoma cell lines against three small molecule inhibitors of ATR (ATRi): AZD-6738/Ceralasertib, VE-822/Berzoserib and BAY-1895344, all of which are currently in early phase clinical evaluation, either alone or in combination with DNA damaging chemotherapies [50–52]. In order to compare the responses, we deduced the area under the curve (AUC) based on nine-point concentration response curves established for each line within the panel. We found that ALT-negative osteosarcoma cell lines with short telomeres were significantly more sensitive to all three of the small molecule ATRi compared to the ALT-negative lines with long telomeres or the ALT-positive lines (Figure 3). We observed no significant difference by TMM status against methotrexate (Supplementary Figure 3), a drug commonly used to treat osteosarcoma that disturbs DNA replication and induce DNA damage through the inhibition of dihydrofolate reductase. Thus, association of the drug sensitivity with short telomeres is selective for ATRi.

**Figure 3.**
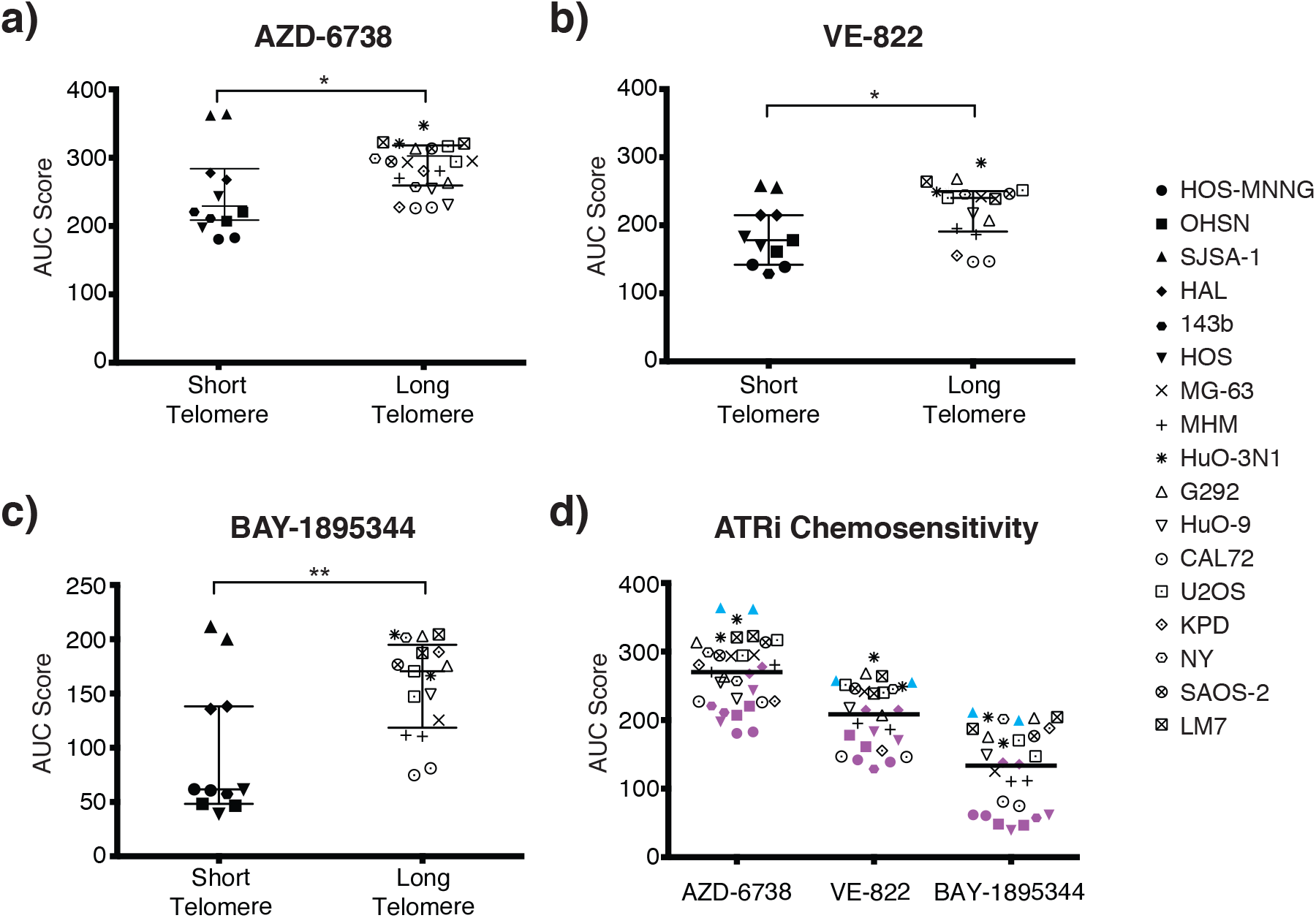
ATRi sensitivity in osteosarcoma cell lines. a-c) Sensitivity of osteosarcoma lines to ATRi VE-822 (a), AZD-6738 (b), BAY-1895344 (c). Data represent AUC deduced from concentration response survival curves. Osteosarcoma lines grouped according to telomere length. Individual lines represented by symbols on the right (short telomere lines n=6, long telomere lines n=11). Values summarise outcome from two repeats for each cell line. Bars indicate median AUC and 95% confidence interval for each group, Mann-Whitney U test * p<0.05, ** p<0.01. d) Summary of ATRi sensitivity. Short telomere cell lines are highlighted in purple. An outlier, ATRi-unresponsive short telomere line, SJSA-1, is marked in cyan.

Among the three ATRi, the most recently developed BAY-1895344 was the most potent, yielding the greatest differential between groups (Figure 3d). The half maximal inhibitory concentration (IC50) also support this notion (Supplementary Table 4). Collectively, our data indicate that ATRi preferentially eliminate osteosarcoma with short telomeres.

### ATRi-induced chromosome segregation failure and cell death in osteosarcoma with short telomeres

To investigate the cause of the increased ATRi sensitivity seen in the osteosarcoma lines with short telomeres, we monitored cell fate in the presence of VE-822 at concentrations of 0.3, 1 and 3 μM using live cell imaging. We collected phase contrast images and, concurrently, scored for the incorporation of the life cell impermeable SYTOX™ DNA-binding dye, which we added to the medium and which permits real time quantification of cell death. This analysis revealed a strong cytopathic response concurrent with cell death, observed within 24-36 hours of ATRi addition in OHSN, HOS-MNNG and HAL, three tested osteosarcoma cell lines with short telomeres (Figure 4a-d). In contrast, MG-63 and MHM, two cell lines with long telomeres, did not respond with death or evidence of cytopathic response within the 48-hour observation period (Figure 4a, e and f). Similar results were obtained when cells were treated with the ATR inhibitor BAY-1895344 (Supplementary Figure 4b-f). Notably, an overt cytopathic response remained observable in short telomere cells where SYTOX™ dye and/or fluorescence imaging to detect dye incorporation was omitted, indicative that the response is not linked to intercalation of the dye into DNA or exposure of the cultures to fluorescent light (Supplemental figure 4a). Thus, our data indicate that ATR activity is required for the survival of osteosarcoma with short telomeres.

**Figure 4.**
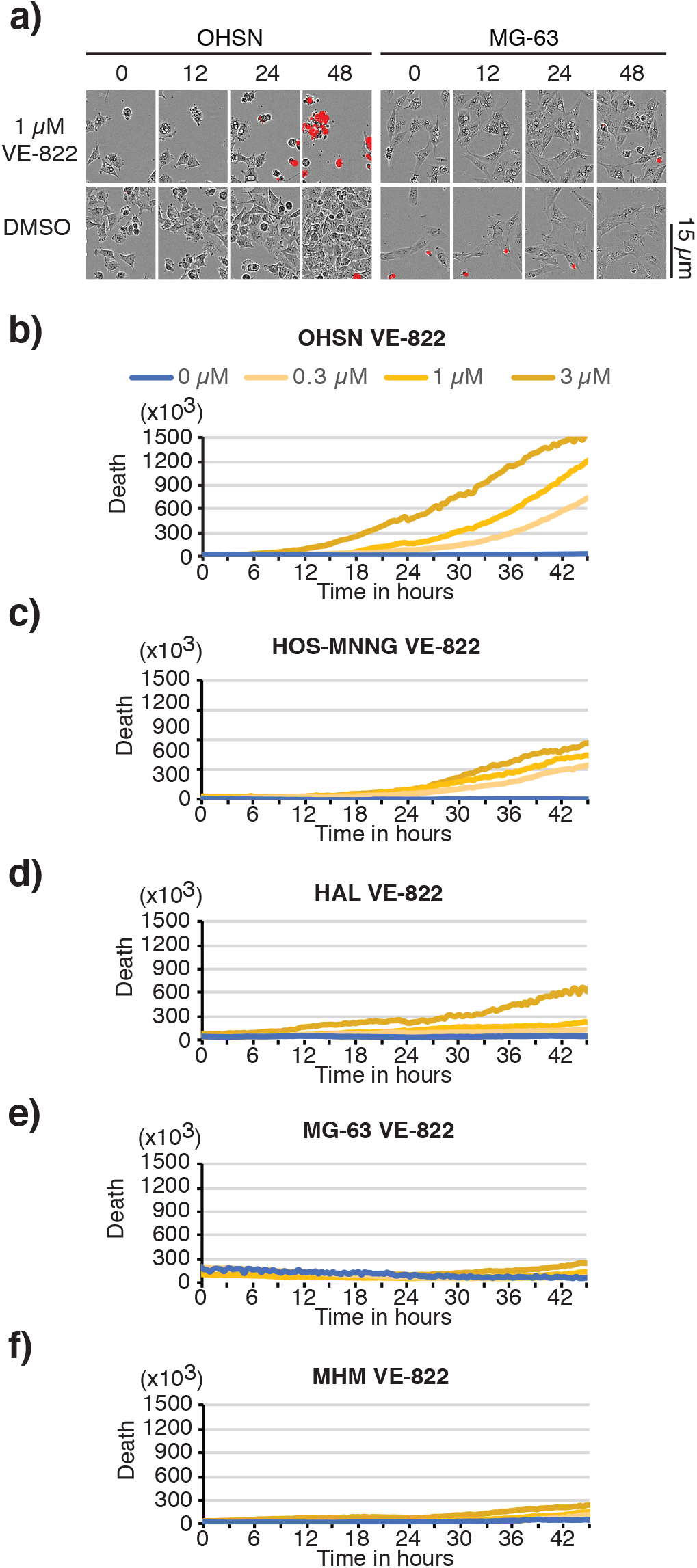
Selective death of osteosarcoma with short telomeres following treatment with ATR inhibitor VE-822. Short telomere cell lines, OHSN, HOS-MNNG and HAL, and long telomere cell lines, MG-63 and MHM, treated with VE-822 were assessed using IncuCyte live cell analysis. Medium was supplemented with the life cell-impermeable dye SYTOX™ to quantify cell death. a) Representative images of OHSN and MG-63 treated with vehicle (DMSO) or 1 μM VE-822 at the indicated time. A 15-μm scale bar is shown. Phase contrast images are overlayed with SYTOX™ death dye signal (red) marking dead cells. b-f) Representative graphs depicting net death over time, reflected by SYTOX™ death dye incorporation normalised to cell density for OHSN (b), HOS-MNNG (c), d) HAL (d), MG-63 (e) and MHM (f). Data representative of two or more experiments.

Lack of telomere replication is expected to lead to loss of telomere protection, in turn eliciting chromosomal fusions and, as a result, defective mitosis with peri- and postmitotic death. To probe for such events, we made use of GFP-fused histone H2B (GFP-H2B), which we expressed in OHSN cells featuring short telomeres and, for comparison, NY cells featuring long telomeres. Longitudinal live-cell-analysis revealed a prominent increase in mitotic DNA bridges detectable as early as 6-hours following ATRi addition in the short telomere OHSN cells but not long telomere NY cells (Figure 5a, cyan arrows). We also observed chromatin fragmentation and apoptotic DNA bodies in OHSN cells but not NY cells (Figure 5a, yellow arrows) with a dramatic increase in prevalence over time (Figure 5b). Thus, we concluded that ATR inhibition causes selective cell death preceded by prominent mitotic aberrations consistent with chromosome fusions in osteosarcoma with short telomeres.

**Figure 5.**
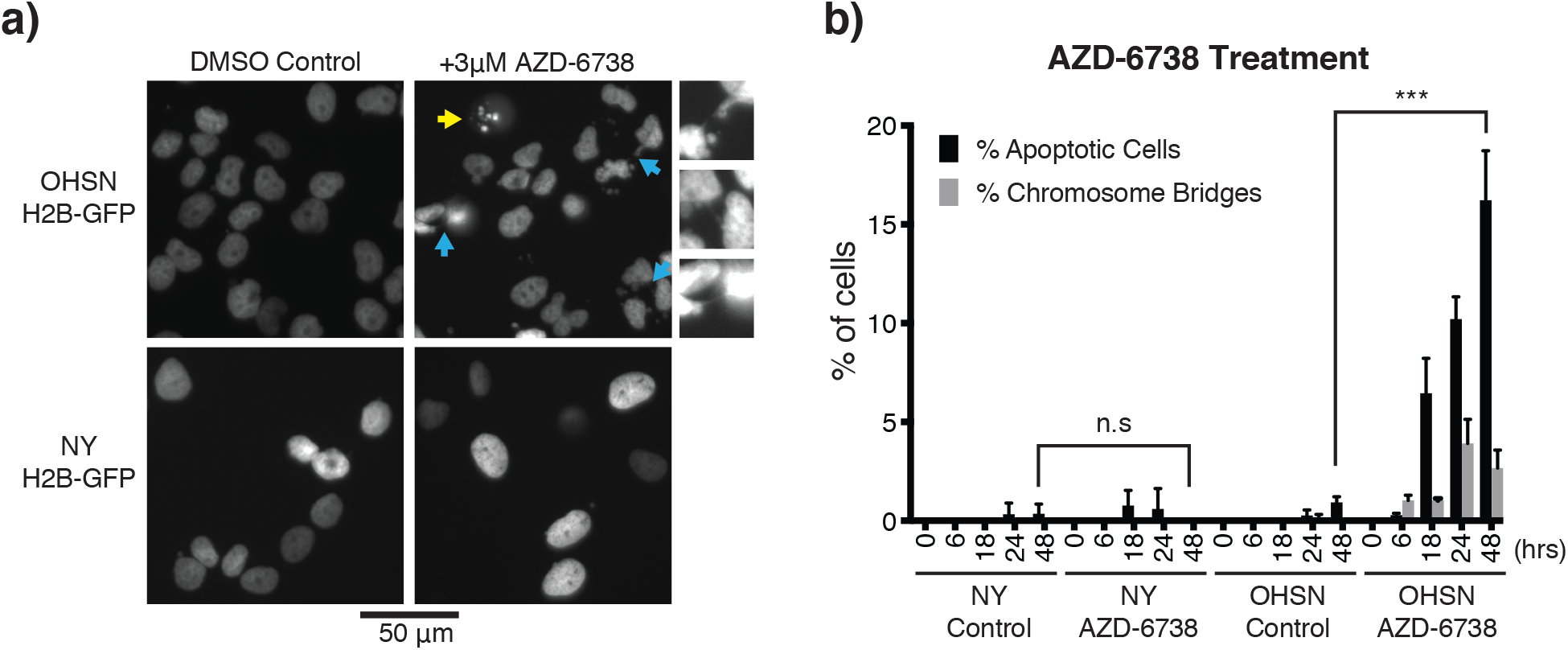
ATRi-induced chromosome bridges and apoptotic cell death in osteosarcoma with short telomeres. a) *Representative images of OHSN and NY cell lines tagged with H2B-GFP.* Yellow arrow shows apoptotic cell. Cyan arrows show chromosome bridges, which are enlarged and shown on the right of the panel. A 50-μm scale bar is shown. b) *Quantification of apoptotic cells and chromosome bridges over 48 hours*. Cells were treated with vehicle or AZD-6738 and were imaged at 6, 18, 24 and 48 hours to count number of apoptotic cells and chromosome bridges from randomly selected samples of at least 200 cells. Graph shows mean percentage of cells classified as apoptotic or containing chromosome bridges for three independent experiments. Error bars show standard deviation. Short telomere cell line OHSN has significantly more apoptotic cells after AZD-6738 treatment compared to control cells, while no significant difference is seen for the long telomere ALT-positive cell line NY (Student T-test, *** p<0.001).

## Discussion

In this study, we demonstrated that the mode of telomere maintenance can influence ATRi sensitivity and addressed the reasons for efficacy of ATRi in the case of osteosarcoma. Our study using a panel of seventeen osteosarcoma-derived cancer cell lines revealed that ALT-negative lines with short telomeres are hypersensitive to ATRi. Although ATR is an essential factor for genome stability and cell survival, our cytological analysis showed that only osteosarcoma cell lines with short telomeres died within 1-2 days in the presence of the ATR inhibitors. Inactivation of ATR resulted in chromosome bridges in mitosis and cell death. Thus, ATRi selectively eliminate osteosarcoma with short telomeres. Our results highlight the possibility of selectively exploiting the mechanisms of short telomere maintenance for cancer therapy.

We report comprehensive parallel characterisation of TMM covering an extensive panel of osteosarcoma derived lines. Our study agrees with the view that the level of telomerase expression does not strongly correlate with the length of telomeres that are maintained. Of note, we describe one cell line, HuO-3N1, which categorised as ALT-negative with long telomeres and had low or undetectable levels of TERT. Such cancers were recently reported [6–8], and HuO-3N1 may reflect a model with this molecular feature. Together, our panel of osteosarcoma cell lines covers distinct TMMs and would be a useful tool to study cancer-specific telomere biology and TMM-based cancer therapeutics.

### The necessity of ATR activity at short telomeres in osteosarcoma

A past report suggested that one ATRi, VE-822, selectively kills ALT-positive cells [44]. Other reports have failed to detect this association [53–56]. These studies focused on ALT status but not telomere length and did not include sufficient numbers of osteosarcoma cell lines that maintain short telomeres, except SJSA-1, which exhibited significant resistance to a wide range of kinase inhibitors in a recent large-scale profiling study [33]. Our data provides evidence for the first time that replication of naturally sub-normally short telomeres, as occurs in human cancers, requires ATR activity. Telomere replication is inherently difficult due to obstacles encountered by the replication fork such as the repetitive nature of the DNA sequence and the propensity to form secondary structures such as G-quadruplexes [57]. A requirement for ATR in telomere replication has been suggested in studies using cell lines derived from patients with Seckel Syndrome who have a hypomorphic mutation in *ATR* (OMIM 210600) [58]. Similar telomere instability has been recaptured in the Seckel syndrome mouse model [19]. Notably, frequent chromosome fusions were observed in the Seckel mouse cells where short telomeres were experimentally induced by deletion of telomerase RNA. This observation is consistent with our finding, documenting elevated chromosome separation defects and cell death in osteosarcoma with short telomeres in the presence of ATR inhibitors.

How short telomeres activate ATR remains to be established. However, various lines of evidence predict a role for ATR in this context. The telomeric DNA binding protein, TRF1, facilitates telomere replication, and reduced TRF1 binding is known to induce telomere replication defects, ATR activation and telomeric ultrafine bridges [59–61]. We speculate that shortened telomeres with reduced TRF1 density regularly activate ATR in order to facilitate telomere replication. ATR, together with the related kinase ataxia telangiectasia mutated (ATM), is thought to have a role in navigating telomerase to the telomere to extend it [62]. Thus, localised ATR activation, mediated by fork-stalling at telomeric sequences, may enhance telomerase recruitment, as emphasised in the replication-mediated telomerase recruitment model [63].

### Telomere diagnostic as a potential tool for cancer stratification

Although Southern blot-based TRF analysis has been a gold standard for telomere length analysis, it is labour-intensive and requires a large quantity of intact genomic DNA, making it unsuitable in a clinical setting. Our MM-qPCR analysis for the panel of osteosarcoma cell lines highly correlated with the TRF results. Although cancer cells can be aneuploid or polyploid, the recent development of MM-qPCR made it possible to estimate relative telomere length of cancer [41, 42]. We were able to clearly identify a group of short telomere-type TMM in our panel using this technique by comparing telomere repeat number in these cells to those in the reference cell line, HEK293T. As this PCR-based assay is simple, rapid, high-throughput and needs only a small quantity of sample for diagnosis [40, 64, 65], we believe this assay is robust and can be used for the primary assessments of telomere repeat number and ALT-status.

In summary, our data demonstrates that short telomeres predict hypersensitivity to ATRi in osteosarcoma. We propose the use of telomere length as a biomarker to predict high efficacy of ATRi.

## Material and Methods

### Cell Lines

Cell lines and media used in this study are listed in Supplementary Table 1. Cells were sourced from the Sanger Centre cell line project [66]. H2B-GFP expressing lines were generated as described in [67]. All cells were maintained at 37°C in a humidified incubator with 5% CO_2_.

### DNA/RNA Extraction

Cell pellets were collected from actively proliferating cells grown in 10-cm plates to 70-80% confluency. Genomic DNA and total cellular RNA were extracted using commercially available kits (Invitrogen).

### Telomere length measurements

Telomere Restriction Fragment (TRF) analysis and monochrome-multiplex quantitative polymerase chain reaction (MM-qPCR) were performed, as described previously [68].

Briefly, TRF assays used 4 μg of genomic DNA digested with *Hin*fI and *Rsa*I and separated on a 30-cm 0.8% agarose gel. Genomic DNA fragments were probed with an alpha-^32^P-labelled telomeric DNA fragment, released using *Eco*RI from pKazu-hTelo, containing 55xTTAGGG repeats, cloned into pCR4 TOPO-blunt vector (Invitrogen). DNA weight marker Hyperladder™ 1-kb (BIoline) and UltraRanger 1kb (Norgen Biotek) were probed with a *S. pombe* telomeric DNA, released from plasmid p1742 [69].

MM-qPCR was carried out as described in [40] with minor alterations. Genomic DNA samples were diluted to 5 ng/μl. Primer sets hTeloG/C (T), AlbuminF/R and GlobinF/R (S) are listed in Supplementary Table 2. Five concentrations of reference genomic DNA purified from HEK293T (ATCC® CRL-3216™) were prepared by 3-fold serial dilution (from 150 ng to 1.85 ng) to generate standard curves for relative quantitation of T/S ratios. 20 ng of genomic DNA was mixed with 0.75x PowerUp SYBR Green Master Mix (Thermo Scientific), the primers (300 nM) and water, to a final volume of 20 μl per well, and analysed using a Thermo Fisher QuantStudio 5, involving denaturation for 15 minutes at 95°C, followed by two cycles of 15 seconds at 94°C, 15 seconds at 49°C and 32 cycles of 15 seconds at 94°C, 10 seconds at 62°C, 15 seconds at 74°C with signal acquisition and 10 seconds at 84°C, 15 seconds at 88°C with signal acquisition. Samples were analysed in triplicate, and analysis was repeated at least three times using independent runs. Data with more than 1% of a coefficient of variance for average Ct were rejected. Telomere copy number was expressed as fold-enrichment against that in the reference HEK293T genomic DNA.

### C-circle assay

CC-assays were carried out as previously described [64, 65]. Briefly, 20 μl CC reactions containing 4 mM dithiothreitol (DTT), 0.1% Tween, 1 mM dNTP mix, 0.2 μg/ml bovine serum albumin (BSA), 7.5U phi-29 polymerase (New England Bioscience), 1x phi-29 buffer and 32 ng of gDNA were incubated at 30°C for 8 hours followed by incubation at 65°C for 20 minutes. For each sample tested, parallel reactions were carried out without addition of phi-29 polymerase.

For MM-qPCR based quantification, 10 μl of the CC-assay products were diluted with 30 μl Tris-EDTA buffer and 5 μl of diluted products were used to quantify the C-circles. Standard curves were generated using serial three-fold dilutions of U2OS DNA, ranging from 3.2 ng/μl to 0.013 ng/μl. qPCR reaction was run in triplicate and contained 5 μl DNA, 1x PowerUp SYBR Green Master Mix, 10 mM DTT, 2% DMSO and 500 nM of the primer sets. Primer sets used were CC-TeloF/R (T) and AlbuminF/R (S). The following PCR conditions was used: 15 minutes at 95°C; 33 cycles of 15 seconds at 95°, 120 seconds at 54°C. Ct values were then extracted for analysis and expressed as T/S. CC assay (CCA) level were calculated by subtracting T/S by that of the respective no phi-29 polymerase controls. A sample is then considered to be ALT-positive if its CCA level lower error bar is greater than zero.

For dot blot quantification, CC-assay products were diluted with 200 μl 2xSSC and transferred to a nitrocellulose membrane using a dot blot vacuum manifold. DNA was UV-cross linked on the membrane and probed using human telomere probe, as described for TRF.

### Protein Extraction and Immunoblotting

Cell were lysed into 2% w/v SDS, 50mM Tris pH6.8. Lysates were separated using discontinuous SDS-PAGE to resolve proteins <200kDa or Tris-Acetate gels (NuPAGE, Invitrogen) to resolve proteins >200kDa. Antibodies used for immunoblotting were anti-SMARCAL1 (1/100, D3P5I; Cell Signaling), anti-DAXX (1/1000, 25C12; Cell Signaling), anti-ATRX (1/1000, D1N2E; Cell Signaling), anti-ATRX (1/100, sc-55584; Santa Cruz Biotechnology), anti-alpha-tubulin (1/5000, DM1A; Sigma Aldrich) and anti-GAPDH (1/5000, ab9485; Abcam).

### RNA quantification by Reverse-transcription quantitative polymerase chain reaction (RT-qPCR)

RNA quantification using RT-qPCR was carried out as described previously [69]. Briefly, 1 μg of total RNA was reverse-transcribed to cDNA using 200U SuperScript IV reverse transcriptase (Thermo Scientific) and 50 μM random hexamer oligos. Reverse transcribed cDNA was diluted to 200 μL with UltraPure DEPC-treated water, remaining RNA was removed with RNase A and samples analysed using PowerUp SYBR Green based real-time PCR. Primers sets used were hTERT F1579/R1616, hTERC F27/R163, 7SK F7/R112, GAPDH F6/R231 (Supplementary Table 2). qPCR cycling conditions were 95°C for 10 minutes followed by 40 cycles of 10 seconds at 95°C, 20 seconds at 60°C and 10 seconds at 72°C. Melting curve analysis was performed immediately thereafter. *TERT* and *TERC* expression was normalised to *7SK* mRNA expression using the ΔCT method. All data are expressed as ΔΔCT compared with HEK293T or as percentage expression compared to the HEK293T cell line. Averages were calculated from three biological replicates.

### Fluorescence microscopy-based detection of ALT-associated promyelocytic leukaemia (PML) bodies (APB)

Cells were grown on ethanol-treated glass coverslips until they reached 50-80% confluency, then fixed with −20°C methanol for 10 minutes. To detect telomeres and PML bodies, Alexa Fluor® conjugated primary antibodies, anti-TRF2 [Alexa Fluor® 488] (1/100, ab198595; Abcam) and anti-PML [Alexa Fluor® 647] (1/50, ab209955; Abcam), were used respectively. Coverslips were mounted into 4ʹ,6-diamidino-2-phenylindole (DAPI) to counterstain the DNA. Imaging was carried out using a DeltaVision Elite (GE Healthcare).

A cell line was considered APB-positive if APBs (a focus of TRF2 localised within a PML body) were detected in more than 10% of nuclei. At least 50 cells were analysed in each of three separate biological samples.

### Cell Viability Assay

Cell viability was assessed using Resazurin reduction. Cells were seeded at densities optimised for each line into 96 well plates. Cells were exposed to nine concentrations of three-fold serially diluted inhibitors 24 hours post seeding. Assays were run in triplicate. Cell viability was assessed 96 hours post inhibitor addition. Area under the curve metrics (AUC) were deduced based on concentration-response curves normalised to vehicle for each line.

### IncuCyte Live Cell Analysis

Time-lapse observations were carried out using an IncuCyte ZOOM™ live-cell analysis system (Essen Bioscience), as described previously [67]. Cells were seeded into 96-well plates 24 hours prior to ATRi addition followed by imaging every two hours at 37 °C with 5% CO_2_. SYTOX TM green death dye (Molecular Probes) was added at the time of ATRi addition in some experiments and monitored using fluorescence image acquisition. Data presented are typical for at least two independent assessments, with conditions assessed in triplicate.

### Statistical Analysis

Microsoft Excel and GraphPad Prism software were used to carry out statistical analysis for qPCR, chemosensitivity experiments and the extraction of AUC values from cell viability assay data. Mann-Whitney U tests were carried out to determine statistical significance in the chemosensitivity data (* at p<0.05, ** at p<0.01 and *** at p<0.001).

## Acknowledgements

KT and LCC were supported by Cancer Research UK (C36439/A12097) and European research council (281722-HRMCB). GZ and CAM were supported by Children with Cancer UK (17-244). CM was supported through a Cancer Research UK studentship (C416/A23233).

