## Supplementary Tables1-4 Figures1-4 for "Selective Elimination of Osteosarcoma Cell Lines with Short Telomeres by ATR Inhibitors"

Supplementary Table 1 – Origin and Characteristics of the Osteosarcoma Cell Lines
Supplementary Table 2 – Oligos used for this study
Supplementary Table 3 – Telomere length measurement from TRF in Figure 1c
Supplementary Table 4 – IC50 drug response in osteosarcoma cell lines
Supplementary Figure 1 – Characterisation of Telomere Status in Osteosarcoma Cell Lines
Supplementary Figure 2 – APB Assay
Supplementary Figure 3 – Sensitivity to methotrexate and telomere length
Supplementary Figure 4 – Selective death of osteosarcoma with short telomeres exposed to ATR inhibitor BAY-1895344

**Supplementary Table 1 – Origin and Characteristics of the Osteosarcoma Cell Lines**

| Cell Line | Media | Age (Years)[a] | Sex[a] | Reference[b] |
|---|---|---|---|---|
| **HOS-MNNG[c]** | RPMI | 13 | Female | ATCC-CRL-1547 |
| **OHSN** | RPMI | 14 | Male | *Fodstad et al. 1986* [1] |
| **SJSA-1** | RPMI | 19 | Male | ATCC-CRL-2098 |
| **HAL** | RPMI | 16 | Male | ExPASy (CVCL_D788) |
| **143b[c]** | DMEM | 13 | Female | ATCC-CRL-8303 |
| **HOS** | EMEM | 13 | Female | ATCC-CRL-1543 |
| **MG-63** | EMEM | 14 | Male | ATCC-CRL-1427 |
| **MHM** | RPMI | 41 | Female | *Kjønniksen et al. 1994* [2] |
| **HuO-3N1** | RPMI | 15 | Female | ExPASy (CVCL_1297) |
| **G292** | McCoy's | 9 | Female | ATCC-CRL-1423 |
| **HuO-9** | RPMI | 13 | Female | ExPASy (CVCL_1298) |
| **CAL72** | DMEM | 10 | Male | ExPASy (CVCL_1113) |
| **U2OS** | McCoy's | 15 | Female | ATCC-HTB-96 |
| **KPD** | RPMI | 7 | Female | *Bruland et al. 1988* [3] |
| **NY** | RPMI | 15 | Male | ExPASy (CVCL_1613) |
| **SAOS-2** | DMEM | 11 | Female | ATCC-HTB-85 |
| **LM7[d]** | DMEM | 11 | Female | *Jia et al. 1999* [4] |

[a] Age and sex of the patient from whom the tumour originates
[b] Identifiers given for cell lines available at the ATCC or ExPASy
[c] metastatic derivatives of HOS
[d] metastatic derivative of SAOS-2

**Supplementary Table 2 – Oligos used for this study**

| Oligo name | Sequence |
|---|---|
| hTeloG | ACACTAAGGTTTGGGTTTGGGTTTGGGTTTGGGTTAGTGT |
| hTeloC | TGTTAGGTATCCCTATCCCTATCCCTATCCCTATCCCTAACA |
| AlbuminF | CGGCGGCGGGCGGCGCGGGCTGGGCGGAAATGCTGCACAGAATCCTTG |
| AlbuminR | GCCCGGCCCGCCGCGCCCGTCCCGCCGGAAAAGCATGGTCGCCTGTT |
| GlobinF | CGGCGGCGGGCGGCGCGGGCTGGGCGGCTTCATCCACGTTCACCTTG |
| GlobinR | GCCCGGCCCGCCGCGCCCGTCCCGCCGGAGGAGAAGTCTGCCGTT |
| CC-TeloF | GGTTTTTGAGGGTGAGGGTGAGGGTGAGGGTGAGGGT |
| CC-TeloR | TCCCGACTATCCCTATCCCTATCCCTATCCCTATCCCTA |
| hTERT F1579 | GCTGACGTGGAAGATGAGCGTGC |
| hTERT R1616 | TCCTCACGCAGACGGTGCTCTG |
| hTERC F27 | GGTGGTGGCCATTTTTTGTC |
| hTERC R163 | GTAGAATGAACGGTGGAAG |
| 7SK F7 | GAGGGCGATCTGGCTGCGACAT |
| 7SK R112 | ACATGGAGCGGTGAGGGAGGAA |
| GAPDH F6 | GAAGGTGAAGGTCGGAGT |
| GAPDH R231 | GAAGATGGTGATGGGATTTC |

**Supplementary Table 3 – Telomere length measurement from TRF in Figure 1c**

| Cell Line | TMM | Mean Length (kb) | Median Length (kb) | Variance ($kb^2$) | Semi-interquartile range (kb) |
|---|---|---|---|---|---|
| **HEK293T** | Control | 6.64 | 5.74 | 16.72 | 3.64 |
| **HOS-MNNG** | ST | 5.21 | 2.98 | 20.44 | 1.86 |
| **OHSN** | ST | 5.02 | 3.22 | 19.12 | 2.02 |
| **SJSA** | ST | 4.83 | 3.21 | 17.93 | 2.12 |
| **HAL** | ST | 4.71 | 2.70 | 22.74 | 1.90 |
| **143b** | ST | 4.42 | 2.96 | 16.21 | 2.02 |
| **HOS** | ST | 4.70 | 3.58 | 16.73 | 2.36 |
| **MG-63** | LT | 6.78 | 5.77 | 12.29 | 4.46 |
| **MHM** | LT | 6.73 | 6.37 | 12.94 | 4.18 |
| **HuO-3N1** | LT | 6.06 | 6.67 | 20.63 | 1.76 |
| **G292** | ALT | 21.54 | 10.69 | 524.47 | 6.36 |
| **HuO-9** | ALT | 17.27 | 9.67 | 348.21 | 5.72 |
| **CAL72** | ALT | 20.24 | 10.24 | 477.35 | 6.22 |
| **U2OS** | ALT | 25.21 | 13.72 | 577.77 | 7.34 |
| **KPD** | ALT | 18.95 | 9.95 | 392.04 | 5.94 |
| **NY** | ALT | 19.19 | 9.89 | 390.53 | 5.84 |
| **SAOS-2** | ALT | 13.81 | 7.61 | 251.98 | 4.62 |
| **LM7** | ALT | 15.56 | 8.20 | 293.43 | 4.90 |

ST – ALT-negative, short telomere
LT – ALT-negative, long telomere
ALT – ALT-positive

**Supplementary Table 4 – IC50 drug response in osteosarcoma cell lines**

| Cell Line | TMM | AZD-6738 μM | VE-822 μM | BEY-1895344 μM | Methotrexate μM |
|---|---|---|---|---|---|
| **HOS-MNNG** | ST | 0.29±0.02 | 0.12 | 0.017±0.001 | 0.04 |
| **OHSN** | ST | 0.66±0.18 | 0.25±0.06 | 0.013±0.001 | 0.20 |
| **SJSA** | ST | 22.68±3.49 | 1.98±0.18 | 0.537±0.128 | 0.30 |
| **HAL** | ST | 2.26±0.42 | 0.69±0.01 | 0.081±0.003 | 0.07 |
| **143b** | ST | 0.69±0.14 | 0.09 | 0.018 | 0.02 |
| **HOS** | ST | 0.83±0.48 | 0.29±0.06 | 0.014±0.005 | 0.32 |
| **MG-63** | LT | 3.97±0.19 | 1.30 | 0.061 | 197.8 |
| **MHM** | LT | 2.20±0.45 | 0.36±0.05 | 0.044±0.002 | 67.87 |
| **HuO-3N1** | LT | 9.56±3.98 | 3.06±1.85 | 0.233±0.154 | 71.85 |
| **G292** | ALT | 3.16±2.66 | 1.30±1.12 | 0.152±0.070 | 69.11 |
| **HuO-9** | ALT | 1.08±0.57 | 0.75 | 0.024 | 0.06 |
| **CAL72** | ALT | 0.80±0.03 | 0.13 | 0.015±0.002 | 44.30 |
| **U2OS** | ALT | 4.91±2.16 | 1.44±0.18 | 0.066±0.016 | 0.03 |
| **KPD** | ALT | 2.01±1.70 | 0.18 | 0.678 | 57.84 |
| **NY** | ALT | 2.54±1.34 | 1.68 | 0.059 | 0.09 |
| **SAOS-2** | ALT | 5.17±1.29 | 1.58 | 0.206 | 0.01 |
| **LM7** | ALT | 9.92±0.71 | 1.80±0.49 | 0.574±0.365 | 0.60 |

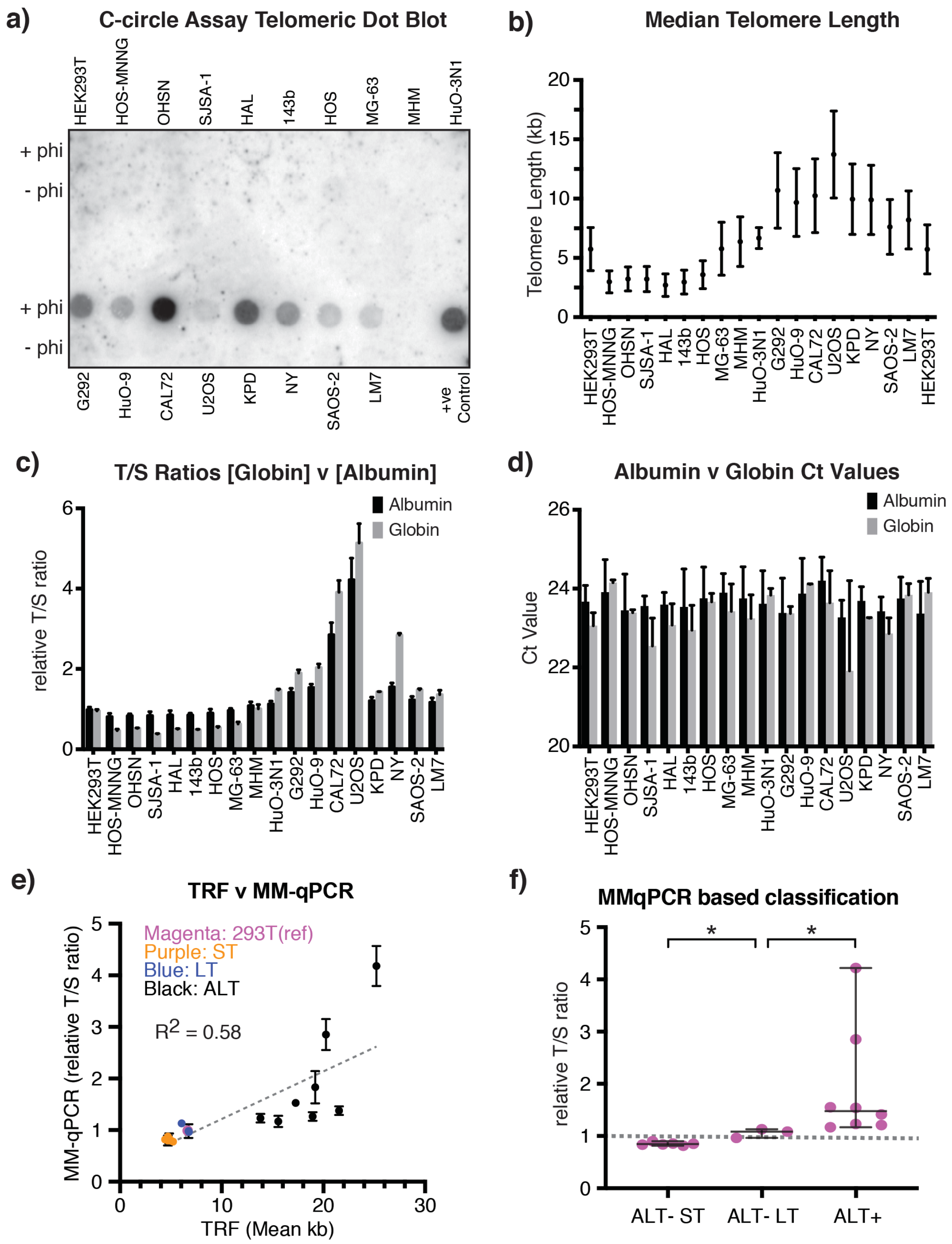


**Supplementary Figure 1 – Characterisation of Telomere Status in Osteosarcoma Cell Lines**

a) *Dot Blot from C-Circle Assay.* C-circle amplification products were detected by dot blot using a telomeric TTAGGG probe. The pCR4 plasmid containing telomeric repeats was used as a positive control. Phi indicates Phi-29 polymerase.

b) *Box plot of median telomere lengths from TRF analysis in Figure 1c as calculated by Telometric software*. Graph shows the median and interquartile range of the telomere fragments.

c) *Comparison of T/S ratios obtained using albumin or beta-globin as the single copy gene (SCG)*. T/S ratio is expressed relative to the T/S ratio determined for HEK293T cell line as expressed as the fold enrichment. Data represent the mean of three independent experiments, each MM-qPCR run in triplicate. Error bars show standard deviation.
d) Uniformity of Ct (cycle threshold) values obtained with *albumin* locus, compared to the *beta-globin* locus. Standard deviation, standard error of the mean and coefficient of variation for the *albumin* and *beta-globin* loci were 0.234 v 0.568, 0.055 v 0.134 and 0.99% v 2.43%, respectively).
e) Correlation graph between TRF-based and MM-qPCR-based telomere length measurements. T/S ratio was determined using *albumin* gene as a SCG, and are expressed relative to that in HEK293T cell line. Liner regression is indicated as grey dashed Line. TMM status of the samples and reference HEK293T are colour coded as indicated.
f) MM-qPCR derived telomere repeats variation in OS cell lines grouped according to TMM status. The grey dashed line indicates the T/S ratio of HEK293T. Mann-Whitney U Test indicates p=0.0238 and p=0.0121 comparing ALT -ST vs ALT-LT and ALT+ vs ALT-LT, respectively.

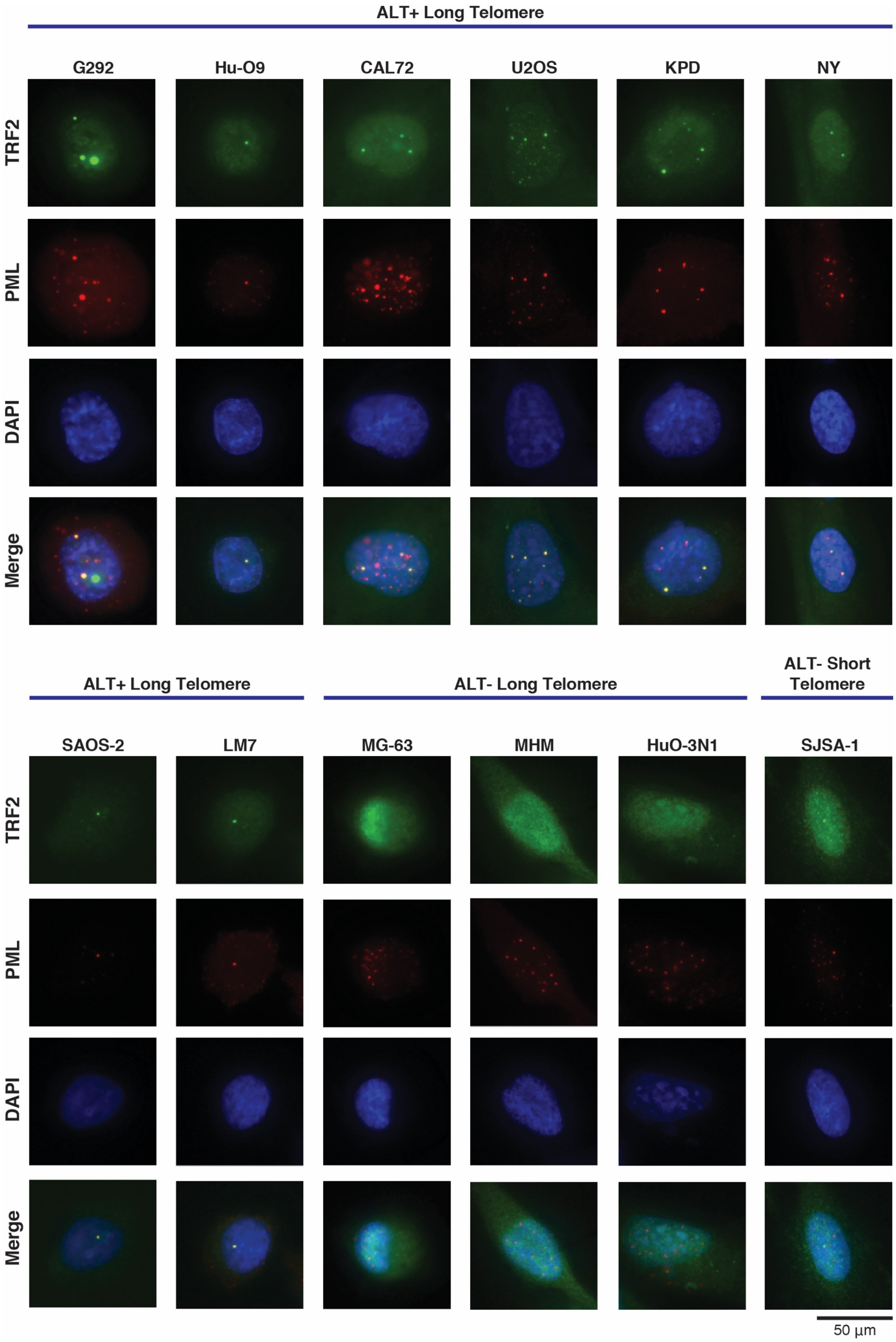


**Supplementary Figure 2 – APB Assay**

Representative immunofluorescence images showing the presence/absence of ALT associated PML bodies (APBs) in the osteosarcoma cell lines with long telomeres. SJSA-1 is shown as a negative control. A 50-µm scale bar is shown.

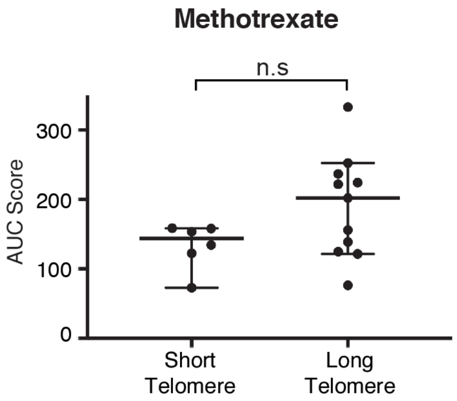


**Supplementary Figure 3 – Sensitivity to methotrexate and telomere length**
Graph shows AUC values deduced for dose response curves to methotrexate. Lines are grouped according to telomere length cell lines. Samples were assessed in triplicate and values averaged. Bars depict median and 95% CI. Mann-Whitney U Test specified p=0.1802, indicating no significant difference between groups.

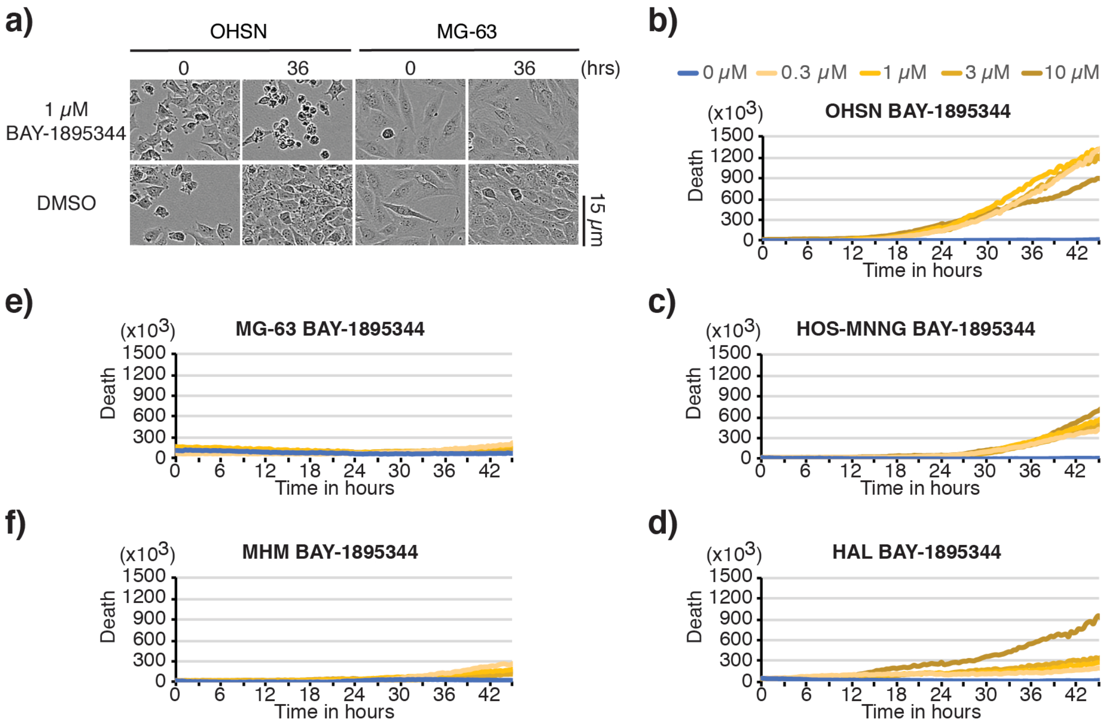


**Supplementary Figure 4 – Selective death of osteosarcoma with short telomeres exposed to ATR inhibitor BAY-1895344**
Short telomere cell lines, OHSN, HOS-MNNG and HAL, and long telomere cell lines, MG-63 and MHM, were treated with VE-822 at the concentration of 0 µM, 0.3 µM, 1 µM and 3 µM and were monitored by *IncuCyte live cell analysis*.
a) Representative images of OHSN and MG-63 with DMSO only and with 1 µM BAY-1895344 at the indicated time are shown. Sample analysis was in the absence of SYTOX™ green death dye and fluorescence imaging. A 15-µm scale bar is shown.
b-f) Representative graphs showing net death over-time for b) OHSN, c) HOS-MNNG, d) HAL, e) MG-63 and f) MHM. Graphs represent one of n=2 independent datasets.
